# Molecular basis of differential parasitism between non-encapsulated and encapsulated *Trichinella* revealed by a high-quality genome assembly

**DOI:** 10.1101/2019.12.17.880336

**Authors:** Xiaolei Liu, Yayan Feng, Xue Bai, Xuelin Wang, Rui Qin, Bin Tang, Xinxin Yu, Yong Yang, Mingyuan Liu, Fei Gao

## Abstract

Understanding roles of repetitive sequences in genomes of parasites could offer insights into their evolution, speciation, and parasitism. As a unique intracellular nematode, *Trichinella* consists of two clades, encapsulated and non-encapsulated. Genomic correlation to the distinct differences between the two clades is still unclear. Here we report an annotated draft reference genome of non-encapsulated *Trichinella*, *T. pseudospiralis*, and performed comparative analyses with encapsulated *T. spiralis*. Genome analysis revealed that, during *Trichinella* evolution, repetitive sequence insertions played an important role in gene family expansion in synergy with DNA methylation, especially for the DNase II members of the phospholipase D superfamily and Glutathione S-transferases. We further identify the genomic and epigenomic regulation of excretory/secretory products in relation to differences in parasitism, pathology and immunology between the two clades *Trichinella*. The present study provided a foundation for further elucidation of mechanism of nurse cell formation and immunoevasion as well as identification of phamarcological and diagnostic targets of trichinellosis.

## Background

Expansion of gene families, especially those that are large or repeatedly involve related process, are thought to play important roles in adaptive evolution (Tsagkogeorga et al. 2017; International Helminth Genomes 2019). Of the process that involved in gene family expansion, an alternative mode mediated by repetitive sequences, including retrotransposons and transposons, have directly or indirectly contribute to the functional evolution of genomes in many ways (Kaessmann 2010; McKenzie and Kronauer 2018). Dissecting the roles roles of repetitive sequences on gene family expansion could be mined for clues about evolutionary impetus events of the genomes. Currently, fewer studies have achieved this goal due to majority of repeat region of the genome were not well assembled, resulting in poor genomic assembly (Wang et al. 2016). Advances in long-read single molecule sequencing technologies have opened new possibilities for the elucidation of complex genomes (Tyson et al. 2018).

The intracellular nematode of the *Trichinella* genus comprises of 12 species and genotypes that parasite a wide range of mammalian hosts (Zarlenga et al. 2006). These parasites could be further categorized of two principal evolutionary clades, the encapsulated represented by *Trichinella spiralis* and non-encapsulated by *Trichinella pseudospiralis*. Both evolutionary clades of *Trichinella spp*. share same life cycles occupying two distinct intracellular niches, intestinal epithelium and skeletal muscle cell. The muscle larvae (ML) of *Trichinella*, released by host gastric fluids, invade intestine epithelium and subsequently develop into adult worms (Ad). The newborn larvae (NBL) delivered by female adults migrate to skeletal muscles and invade into muscle cells where they develop into ML and survive for years (Gottstein et al. 2009). Despite the similarity in life cycles, the two clades showed substantial differences in parasitological, pathological and immunological characteristics (Wu et al. 2001b; Z et al. 2002; Boonmars et al. 2005). These differences were driven by post-miocene expansion, colonization, and host switching, resulting in abundant and heterogeneous repetitive elements in *Trichinella* DNA, according to previous molecular and biochemical studies (DS and G 2000; Zarlenga et al. 2006). Based on this genetic variability, researchers were able to develop radiolabeled probes to differentiate *Trichinella* genotypes (AE et al. 1986; MR et al. 1994). Therefore, the *Trichinella spp*. represent an ideal model of studying the association of repetitive sequences with adaptation to different host ranges. However, further investigation from genome-wide scale is hindered as high quality reference genome was only available for *T. spiralis*, whereas most repetitive sequences were not assembled for the references genomes of the rest *Trichinella spp*. (Mitreva et al. 2011; Korhonen et al. 2016).

To test whether repetitive sequences involved in differential parasitism between the two representative species, we conduct comprehensively comparative genomic and epigenomic analysis between *T. pseudospiralis* and *T. spiralis*, based on a high-quality long-read-assembled reference genome of *T. pseudospiralis*, in which the majority of the repeat regions have now been assembled. We revealed both transposon density and DNA methylation play an important role in parasitic-related gene family expansion. Furthermore, we identified sets of genes related to parasitism whose DNA methylation and expression level varied significantly between the two species. Our work here demonstrated that transposons and methylation contribute to shaping the genome stability and their adaptation to environment via silencing of transposon elements and epigenetic regulation of gene transcription.

## Result

### Genome features

We used one non-encapsulated strain (ISS13) for *de novo* assembly of *T. pseudospiralis* reference genome, using a ‘hybrid’ approach that combined assembly of Pacbio and Illumina reads (Supplementary Tables 1 and 2). After error correction, the total length of the assembled genome was 68.90 Mb, which represents 98.7% of the genome size as estimated by *k*-mer depth distribution of the sequenced Illumina reads (69.79 Mb; Supplementary Fig. 1). The assembly consists of 2,746 scaffolds (≥500 bp) with a mean GC-content of 31.4% and N50 lengths of 208.90 Kb, among which 68 of the largest scaffolds spanning more than half of the *T. pseudospiralis* genome (Table 1). The quality of assembly was further confirmed using the available RNA-seq and EST (expressed sequence tag) sequences (Supplementary Tables 3 and 4). Then we used BUSCO (Benchmarking Universal Single-Copy Orthologs) to assess the completeness of the genome based on the presence of single-copy orthologs from the OrthoDB database. We found that 870 (88.6%) out of 982 genes (nematoda_odb9 lineage) were present and complete in the present study, which are comparable with that of the well-assembled *T. spiralis* genomes. Thus, this assembly was used for further analyses (hereafter named T4_ISS13_R, in comparison with the previously published version “T4_ISS13_r1.0”).

**Table 1.** Summary of Genome Assembly and Annotation for *T. pseudospiralis* and *T. spiralis*.

| Description | *T. spiralis* | *T. pseudospiralis* |
| --- | --- | --- |
| Total scaffolded assembly size (Mb); total scaffolds | 63.53; 6,863 | 68.90; 2,746 |
| Total scaffolds of > 2kb: length (Mb); no. of scaffolds | 57.27; 964 | 68.71; 2,631 |
| Largest scaffold (Mb) | 12.04 | 3.73 |
| N50 in kb of scaffolds; count > N50 length | 6373.44; 4 | 208.90; 65 |
| N90 in kb of scaffolds; count > N90 length | 2.05; 919 | 7.68; 1,231 |
| GC content of the whole genome (%) | 33.9 | 31.49 |
| Number of gene models | 16,186 | 12,682 |
| Mean/Median gene size (bp) | 1837.73/1096 | 2330.43/1077 |
| Mean number of exons per gene | 5.41 | 5.52 |
| Mean/Median exon size (bp) | 178.45/129 | 188.37/137 |
| Repetitive sequences (%) | 20.66 | 40.28 |
| CEGMA (Complete/Partial %) | 94.35/95.56 | 95.56/96.37 |

We predicted 12,682 protein-coding genes, spanning 19.1% of the T4_ISS13_R genome (Supplementary Tables 5). The mean length of all the predicted genes is 2.33 kb, with an average of 5.5 exons and a mean exon length of 188.4 bp per gene (Table 1). Approx. 89.17% of the genes had either known homologs or could be functionally classified (Supplementary Table 6). In addition, we also annotated potential parasitism-related functional proteins, such as proteinases, protein kinases, GPCRs, and excretory/secretory (E/S) proteins, as well as the candidate molecular targets for treatment of trichinellosis. To this end, we have identified 1627 proteinases, 88 GPCRs, 336 kinases, 471 E/S proteins and 154 potential drug targets in *T. pseudospiralis*. A similar number of functional proteins and candidate drug targets were also annotated in the *T. spiralis* genome (Supplementary Data 1). Most of these proteins had support of gene expression data, and could act as important factors in immuno-evasion and excystment/encystment, as suggested by previous studies (Nagano et al. 2009).

In the newly generated version of *T. pseudospiralis* reference genome, a total of 27.76 Mb of non-redundant repetitive elements were then identified, which represents ~40.28% of the genome. Compared to the previously generated *T. pseudospiralis* reference genome, the T4_ISS13_R had approx. 18.9 Mb more repeat sequence than the T4_ISS13_r1.0, accounting for most of the differences in genome size between the two assembly version (Supplementary Table 7).

### Methylome annotation

Previously we confirmed the existence of DNA methylation in the *T. spiralis* genome and its relevance to parasitism (Gao et al. 2012). To further annotate the genome regulation of *T. pseudospiralis*, we applied both bioinformatics analyses of homologous gene sequences and enzymatic tests to confirm the existence and activity of *DNA-methyltransferase 1* (Dnmt1) and Dnmt3 in *T. pseudospiralis* (Fig. 1a; Supplementary Fig. 2). Furthermore, single-base resolution maps of DNA methylation, for three life stages, were generated using whole-genome bisulfite sequencing (Supplementary Figs 3 and 4; Supplementary Table 8). Similar to the *T. spiralis* genome, as previously reported (Gao et al. 2012), the *T. pseudospiralis* genome also displayed stage-specific methylome patterns, in which NBL stages showed rare traces of DNA methylation, while ML and Ad stages were moderately methylated, both for genic regions and repetitive sequences, across the entire genome (Fig. 1b). Between the two genomes, a clear genome-wide methylation difference of non-repetitive regions was observed, as indicated by hierarchical clustering analysis (Fig. 1c). More specifically, *T. pseudospiralis* presented a heavier global methylation level than *T. spiralis* in the non-repeat regions (Fig. 1d). Such significantly higher methylation within the *T. pseudospiralis* genome was not limited to specific regions, but was broadly distributed in genic and intergenic regions across the genome, except for the intron regions (Supplementary Fig. 5). Furthermore, in accordance with a higher level of repeats in the *T. pseudospiralis* genome, a significantly higher methylation level of different repetitive sequences was also observed in the *T. pseudospiralis* genome (Supplementary Fig. 6). These results are in accordance with the CpG content (CpG observed/expected [o/e]) for a total of 1,436 pairs of ortholog single genes being lower in *T. pseudospiralis* than in *T. spiralis*, both across the genome and coding regions (Supplementary Fig. 7). The depletion of normalized CpG o/e values may represent an evolutionary signature of DNA methylation, in animal genomes, as methylated cytosines undergo spontaneous deamination to thymine with high frequency (Sarda et al. 2012).

**Figure 1.**
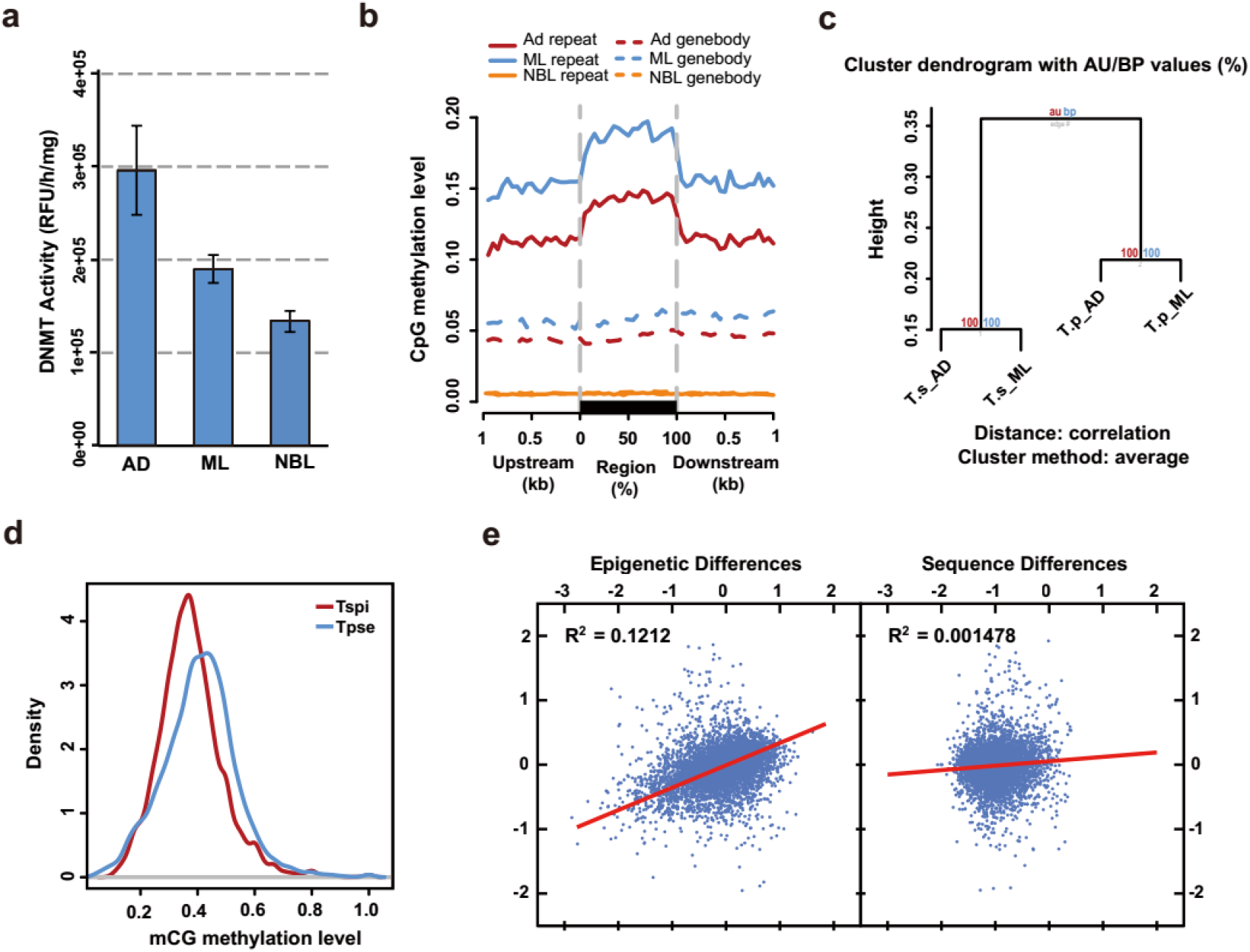
Confirmation and characterization of *T. pseudospiralis* methylome in comparison with *T. spiralis*. **(a)** Results of catalytic activity of *T. pseudospiralis* DNMTs. Triplicates of DNMT activity experiments were carried out, and mean ± SD is indicated. DNMT activity (OD/h/mg) = (Sample OD−Blank OD) / (Protein amount (ug)×hour). **(b)** CpG methylation levels of repeat and gene body regions. Two-kilobase region upstream and downstream of each gene was divided into 100-bp (bp) intervals. Each repeat or gene body region was divided into 20 intervals (5% per interval). **(c)** Clustering of methylation levels of common CpG sites in the whole genome of all the four samples was used in the “Pvclust” algorithm. **(d)** Comparison of mCG methylation level density between *T. pseudospiralis* and *T. spiralis*. **(e)** Correlations among evolutionary changes of epi-modification intensities, gene expression levels, and genomic sequences.

Furthermore, we quantified the epigenetic conservation level using the p-value generated by comparing the common mCGs of orthologous genes between the two species. However, no clear correlation was observed between the methylation and the gene sequence differences (Fig. 1e). However, a much higher correlation was observed between epigenetic and transcriptional divergences (Fig. 1e). These results agree with previous studies that epigenomic conservation is not a simple consequence of sequence similarity, but rather a regulatory mechanism for transcription (Xiao et al. 2012).

### Comparative genomics revealed essential roles of TE and DNA methylation in genome evolution of *Trichinella*

We next evaluated the organizational characteristics of the genomes of *T. pseudospiralis* and *T. spiralis*. The number (12,682) of predicted genes in *T. pseudospiralis* is notably lower than the 16,186 genes identified in *T. spiralis*, which gave a higher gene density in *T. spiralis* (254 per Mb in *T. spiralis* and 184 per Mb in *T. pseudospiralis*; P < 2.2e-16), even though the two genomes showed a similar genome size (63.53 Mb in *T. spiralis* and 68.90 Mb in *T. pseudospiralis*). A comparison of 6857 orthologous genes (based on reciprocal best BLAST hits) in the *T. pseudospiralis* and *T. spiralis* genomes indicated that *T. pseudospiralis* had a significantly longer average intron size compared to *T. spiralis* (240 bp compared to 177 bp; P < 2.2e-16), whereas the average exon size was relatively similar for the two species (199 bp for *T. pseudospiralis* and 194 bp for *T. spiralis*; P = 0.08). A clear correlation was observed between the length of repeats in the introns and intron length, *per se*, in *T. pseudospiralis*, thus indicating that repeats contributed to the greater intron length of *T. pseudospiralis* compared to that in the *T. spiralis* genome (Supplementary Fig. 8). As the most significant difference between the two genomes lay within the repeat content, we next focused on the repeat level between the two genomes. In particular, we noted that within the intergenic regions in *T. pseudospiralis*, 24.1 Mb of repetitive sequences were identified, comprising 86.70% of the total genomic repeats. By contrast, we identified only 8.73 Mb of repeats (63.36 %) within the intergenic regions of the *T. spiralis* genome. Among the various categories, the *T. pseudospiralis* genome contained significantly elevated levels of both tandem repeats and LTRs compared to the *T. spiralis* genome. We identified approx. 11.51 Mb of tandem repeat in *T. pseudospiralis*, based on the TRF method, whereas only 717.76 kb in *T. spiralis*. LTRs were the most dominant type, accounting for approx. 22.23% (15.31 Mb) of the *T. pseudospiralis* genome, much larger than that in the *T. spiralis* genome (4.33 Mb in total) (Supplementary Table 7). After calculating for insertion times, we discovered that a burst of LTR activity likely occurred during the last five million years (Supplementary Fig. 9). Based on these findings, we deduced that these LTRs were most likely inserted into the genome after the divergence of the encapsulated and non-encapsulated *Trichinella*, an event estimated to have occurred 22.6 (15.3 ~ 28.1) million years ago (Supplementary Fig. 11). In addition, the percentage of *de novo* predicted repeats was notably higher than that obtained from homologous predictions, based on Repbase, indicating that *T. pseudospiralis* has many unique repeats compared to other sequenced genomes (Supplementary Table 7).

A Markov clustering algorithm was adapted to compare gene families within phylogenetically-related organisms, in order to delineate gene family expansion and contraction events between the two species. A total of 14,749 orthologous gene families were generated from five nematode species, including two *Trichinella* species (*T. spiralis* and *T. pseudospiralis*), one non-parasite *C. elegans*, one plant parasite, *M. incognita*, one animal parasite, *B. malayi*, and one non-nematode species, *D. melanogaster* (outgroup). Based on the comparison of orthologous gene families, the *T. pseudospiralis* genome displayed 628 expanded and 1,087 contracted gene families, compared with the *Trichinella* common ancestor. By contrast, the *T. spiralis* genome displayed more expanded (1,376) than contracted (884) ones (Fig. 2a; Supplementary Data 2). Among the five species of nematodes, 2,543 families are broadly conserved (NOG), whereas 337 families (containing 1,818 genes) or 684 families (containing 2,577 genes) appear to be *T. pseudospiralis* or *T. spiralis* specific, respectively (Fig. 2b).

**Figure 2.**
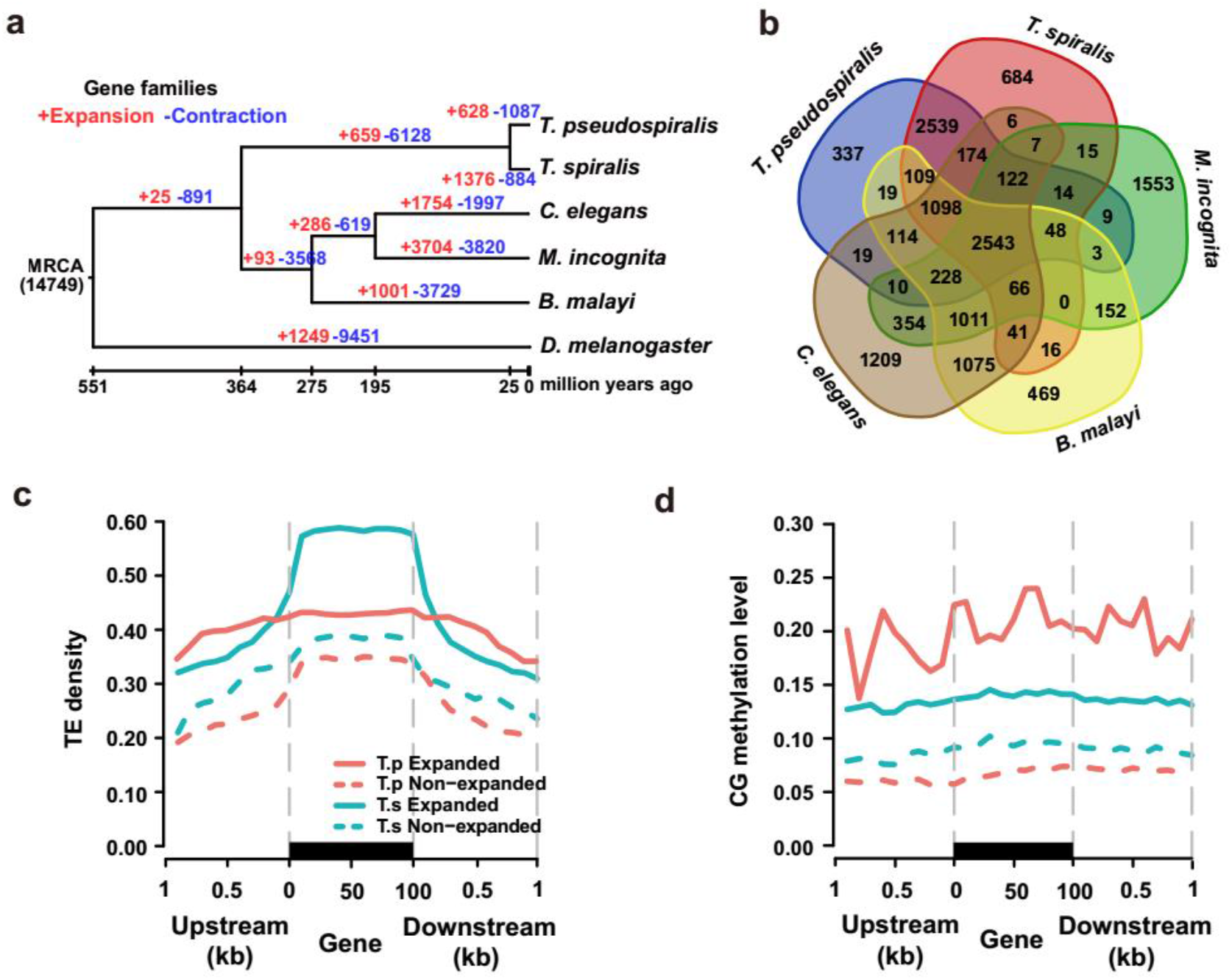
Comparative genomics between *T. pseudospiralis* and *T. spiralis*. **(a)** The dynamics of gene family sizes in the genomes of parasitic nematode *T. pseudospiralis*, *T. spiralis*, *C. elegans*, *M. incognita*, *B. malayi*, with *D. melanogaster* serving as the outgroup. Numbers above the branches represent gene family expansion and contraction events, respectively. Plus sign represents expansion events and the minus sign indicates contraction events. **(b)** Number of shared gene families between *T. pseudospiralis* and four other nematodes (that is, *T. spiralis*, *C. elegans*, *M. incognita* and *B. malayi*). **(c)** Comparison of average density of transposon element (y-axis) between expanded and non-expanded gene families containing TEs around genes and their flanking regions (x-axis). Two-kilobase regions upstream and downstream of each gene were divided into 20 intervals, and so were genic regions. **(d)** Comparison of methylation levels (y-axis) between expanded and non-expanded gene families containing TEs around gene and their flanking regions (x-axis). Two-kilobase regions upstream and downstream of each gene were divided into 20 intervals, and so were genic regions.

Previous studies demonstrated that transposon elements (TEs) are important drivers of gene duplication in adaptation and evolution (Oliver and Greene 2009; Schrader et al. 2014). Considering the big divergence of repeat contents and gene family, we further assessed the TE density with regards to expansion of gene family. A significantly higher level of TE density was observed in the expanded gene families than the non-expanded ones, both in *T. pseudospiralis* (P < 2.2e-16) and *T. spiralis* (P < 2.2e-16) (Fig. 2c). Accordingly, we also observed a significant elevation of DNA methylation levels in the expanded gene families (Fig. 2d), as TEs are highly repressed by DNA methylation (Borgognone et al. 2018). Taken together, these results might suggest that the genome and epigenome divergence between the two clades of *Trichinella* was mainly driven by TEs.

### Transposons facilitate expansion of gene family in synergy with DNA methylation in *Trichinella spp*

To further dissect the potential role of TEs and DNA methylation in the expansion of gene families, we then assessed TE-enriched gene families that were significantly expanded in one species relative to the other (P < 0.05). As a result, we observed 67 and 183 significantly expanded gene families in *T. pseudospiralis* and *T. spiralis*, respectively. Most of these gene families contained novel domains without known function annotations, while only a small number of gene families contained unitary functional domain based on IPR annotations (n=4 in *T. pseudospiralis* and n= 14 in *T. spiralis*, supplementary Table 9).

Among these gene families, the most diversified gene family with known parasitism-related function was the *DNaseII* family. In comparison with *T. pseudospiralis* (19 families, 43 genes), *T. spiralis* displayed significantly more expansion of *DNase II* genes (26 families, 133 genes), representing a unique target for deciphering the mechanism of gene family expansion in *Trichinella* (Supplementary Fig. 10). By reconciling the species phylogeny with the gene phylogeny and by genomic/scaffold location of *DNaseII* genes, we observed that only part, but not all of the family members underwent an expansion (Fig. 3a; Supplementary Fig. 11). More specifically, 8 families containing 15 genes in the *T. pseudospiralis* genome were expanded into 57 genes in the *T. spiralis* genome, but no *T. spiralis* gene families were expanded in *T. pseudospiralis* (Fig. 3b; Supplementary Table 10). Intriguingly, we determined that a majority of these expanded genes were derived from tandem duplication (59.64% in *T. spiralis* and 66.67% in *T. pseudospiralis*), along with some segmental and dispersed duplication events, compared with whole-genome duplication pattern (Supplementary Figs 12 and 13). In order to gain mechanistic insights into the *DNaseII* repertoire expansion, we investigate the repeated context of *DNaseII* superfamily. The differential retentions of duplicated *DNaseII* genes in *T. spiralis* might indicate multiple functions through neo-/sub-functionalization with significantly divergent sequences due to the divergent repeat sequence insertion (Dermitzakis and Clark 2001) (Fig. 3d), as most duplicated *DNaseII* loci were novel repeat families (Supplementary Fig. 14). Despite of that, we also observed that the transposon density of *DNaseII* loci in *T. spiralis* is approximately 1.5× higher than that in the entire genome (Supplementary Fig. 15), meanwhile significantly higher than that in *T. pseudospiralis* (Fig. 3c). This result is consistent with a previous study that has demonstrated high-level transposon density might potentially facilitate *DNaseII* adjacent duplication by non-homologous end-joining along with *Trichinella* evolution history (McKenzie and Kronauer 2018). Next, we tested whether the gene expression reduction occurred in response to gene duplication. As a result, the Spearman correlation coefficients (*r*) between ΔE_Ts − Tp_ and S_Ts/Tp_ were −0.74 and −0.23 in Ad and ML stages, respectively, confirming the prevalence of post-duplication dosage rebalance. Accordingly, we observed significantly higher methylation level in expanded genes than that of non-expanded genes in gene-body and promoter regions both in Ad and ML stages (Fig. 3e; Supplementary Fig. 16), resulting in higher expression level in the non-expanded genes than that of expanded genes both in Ad and ML stages (Fig. 3f).

**Figure 3.**
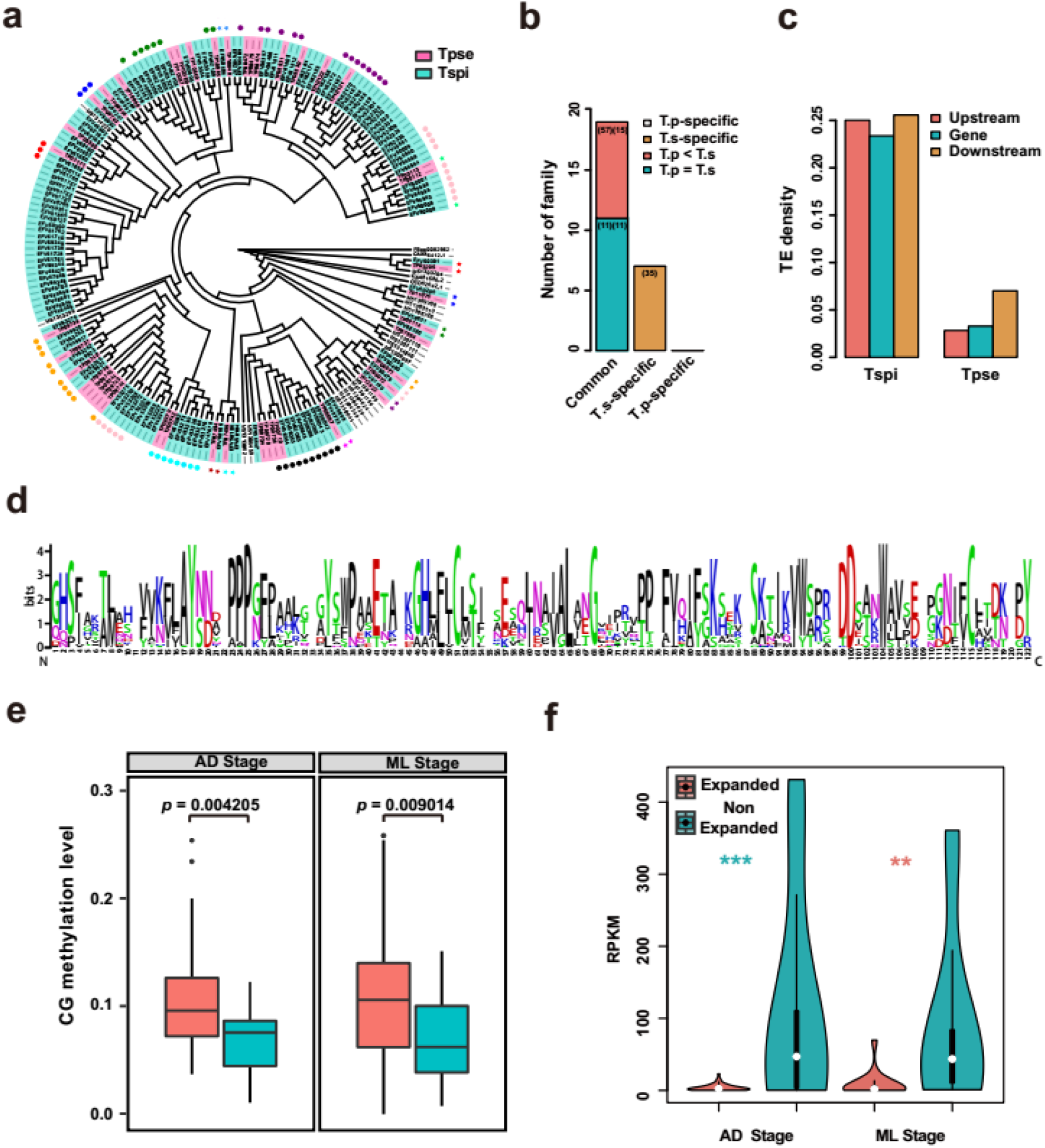
Transposons facilitate duplication of *DNaseII* gene family in synergy with DNA methylation in *T. spiralis*. **(a)** Phylogenetic tree of *DnaseII* gene family with *T. pseudospiralis* (colored in pink), *T. spiralis* (colored in cyan) and other three species (that is *D. melanogaster*, *C. elegans* and *T. suis*). **(b)** Common and specific number of *DnaseII* family between *T. pseudospiralis* and *T. spiralis*. Numbers in the front and back brackets represent gene number and family number, respectively. **(c)** Comparison of average density of transposon element (y-axis) around *DNaseII* loci and their two-kilobase flanking regions upstream and downstream in *T. pseudospiralis* and *T. spiralis* (x-axis). **(d)** Sequence logo of expanded *DNaseII* genes shows the conserved and distinct sequence characteristics in *T. spiralis.* The sequence logo was generated by WebLogo (http://weblogo.threeplusone.com/) from the expanded 57 *DNaseII* sequences aligned at conserved blocks selected by Gblocks. **(e)** Comparison of methylation level between expanded and non-expanded *DNaseII* genes in Ad (left) and ML (right) stages in *T. spiralis*. **(f)** Comparison of expression level between expanded and non-expanded *DNaseII* genes in Ad and ML stages in *T. spiralis*. A two-tailed Student’s test was applied to the pairwise comparison. *** indicates p < 0.001; ** indicates p < 0.01.

In contrast, *T. pseudospiralis* showed significant more expansion of the TB2/DP1/HVA22-related protein (IPR004345) family (n=61 in *T. pseudospiralis* versus n=18 in *T. spiralis*) and the Glutathione S-transferases (IPR004046) family (n=13 versus n=7) (Supplementary Table 9), both of which were reported to be related with parasitism previously. Especially, previous studies showed that Glutathione S-transferases (GST) was likely related with the larval invasion and migration into intestinal epithelial cells (IECs) in *T. spiralis*, and vaccination with GSTs induced a low protective immunity against *T. spiralis* infection (Li et al. 2015; Liu et al. 2018). In *T. pseudospiralis*, phylogenetic analysis revealed that apart from the known GSTs that showed homologous with GSTs in *T. spiralis* (TP07287 and TP02356), *T. pseudospiralis* specifically expanded one gene family (Fam1884), which contained 8 genes (Fig. 4a). Most of these genes were well assembled based on high coverage folds (90× ~ 150×) and had an average of 9 potential antigenic epitopes predicted by ABCpred (artificial neural network based on B-cell epitope prediction server), implying that this expansion played an important role in *T. pseudospiralis* invasion and migration in order to adapt to its non-encapsulated phenotype. Different from the *DNaseII* genes in *T. spiralis*, a majority of the expanded GST genes in *T. pseudospiralis* were derived from dispersed and proximal duplication, with significantly higher TE density observed in the upstream and downstream regions of GSTs (Fig. 4b), potentially facilitating interspersed duplication by nonallelic homologous recombination via surrounding repetitive sequence, according to previous studies (Mendivil Ramos and Ferrier 2012). Likewise, DNA methylation also involved in the dosage rebalance of duplicated GSTs in *T. pseudospiralis*, as higher methylation levels and lower average expression levels were observed in *T. pseudospiralis* (Figs 4c and 4d). Collectively, these results suggest that enriched transposons may potentially facilitate segmental duplication, in synergy with DNA methylation in *Trichinella*.

**Figure 4.**
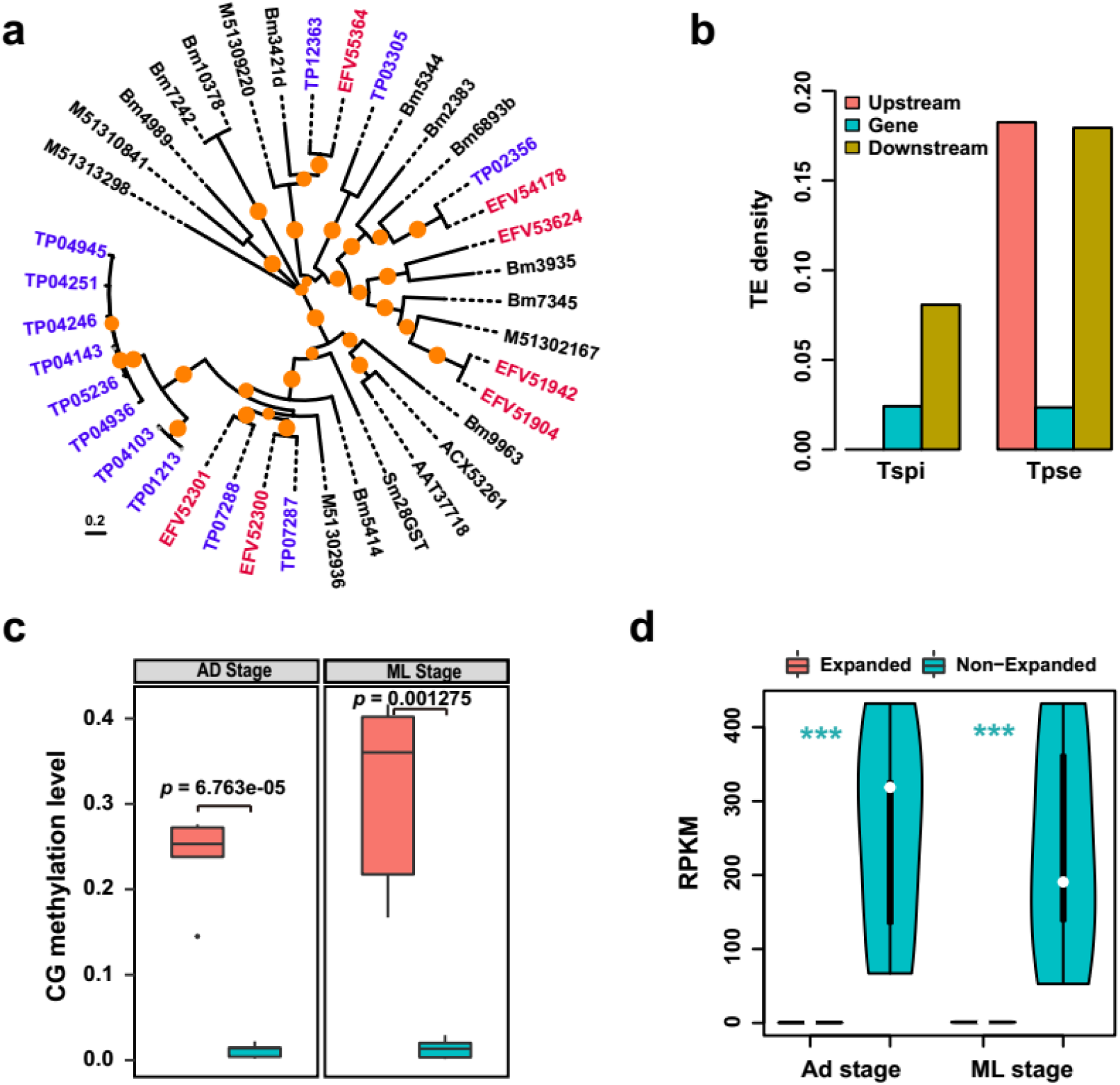
Transposons facilitate duplication of Glutathione S-transferases family in synergy with DNA methylation in *T. pseudospiralis*. **(a)** Phylogenetic relationship of GSTs in *T. pseudospiralis*, other nematodes (that is *T. spiralis*, *B. malayi* and *T. sui*) and GSTs with experimentally verified (that is Sm28GST, ACX53261, AAT37718, EFV54178 and EFV52300). Gene names in blue are from *T. pseudospiralis* and those in red are from *T. spiralis*. Nodes with > 30% bootstrap support (1000 replicates) are indicated in orange circles. **(b)** Comparison of average density of transposon element (y-axis) around GSTs and their two-kilobase flanking regions upstream and downstream in *T. pseudospiralis* and *T. spiralis* (x-axis). **(c)** Comparison of methylation level between expanded and non-expanded GSTs in Ad (left) and ML (right) stages in *T. pseudospiralis*. **(d)** Comparison of expression level between expanded and non-expanded GSTs in Ad and ML stages in *T. pseudospiralis*. A two-tailed Student’s test was applied to the pairwise comparison. *** indicates p < 0.001.

### Divergent E/S genes in relation with varied parasitism in *Trichinella*

The E/S proteins are central to understand parasite-host interactions or host cell modification (Robinson et al. 2007). Among the 471 E/S genes we identified in *T. pseudospiralis*, a large proportion were divergent gene families from *T. spiralis*, including *DNase II*, and TB2/DP1/HVA22-related protein that contained high-level TEs (Supplementary Data 3), again suggesting TEs and DNA methylation play important roles in the differential parasitism of *Trichinella*. On the other hand, the non-TE-containing secreted C-type lectin gene family also had undergone significant expansion in *T. pseudospiralis*, as nine C-type lectins (or lectin-like) E/S genes were identified in *T. pseudospiralis*, but only a single gene was identified in *T. spiralis*. Seven of these lectin genes were supported by RNA-seq data in *T. pseudospiralis*, thereby providing a significantly richer expression pool compared to *T. spiralis* (Fig. 5a). Six of the *T. pseudospiralis* lectins and the one *T. spiralis* lectin showed more homology to *C. elegans* lectin genes, whereas the other three *T. pseudospiralis* lectins shared a greater level of identity with mammalian C-type lectin domains (Fig. 5a), which may play crucial roles for the host-parasite interface, notably in immune evasion, according to previous studies (Loukas 2000). Sequence alignment of the three proteins with their mammalian lectin homologs indicated that the key cysteine residues were perfectly conserved in *T. pseudospiralis* lectin proteins, along with additional residues implicated in forming the stable hydrophobic protein core (Supplementary Fig. 17; Supplementary Table 11). Structure modeling predicted that TP12499 displayed a remarkable structural similarity to the mammalian immune-system lectin Mincle in the PDB library, which can induce acquired immunity such as antigen-specific T-cell responses and antibody production (Lu et al. 2018), and thus, it may well form stable complexes with galactose, as suggested by the high C-score of the models (C-score=0.46). In addition, TP02221 exhibited high structural similarity with MRC2 and, hence, may also bind with mannopyranoside (C-score=0.44) to implement cell infection through endocytosis and this then contributes to antigen presentation (Al-Mulla Hummadi et al. 2006) (Fig. 5b). Finally, TP10242 shared a similar structure with Dectin-2 and may, therefore, function through a raffinose binding site (C-score=0.17) (Supplementary Fig. 18; Supplementary Table 11). Thereby, through these expanded C-type lectin superfamily members, *T. pseudospiralis* might have developed an enhanced capacity for immune evasion compared to *T. spiralis*.

**Figure 5.**
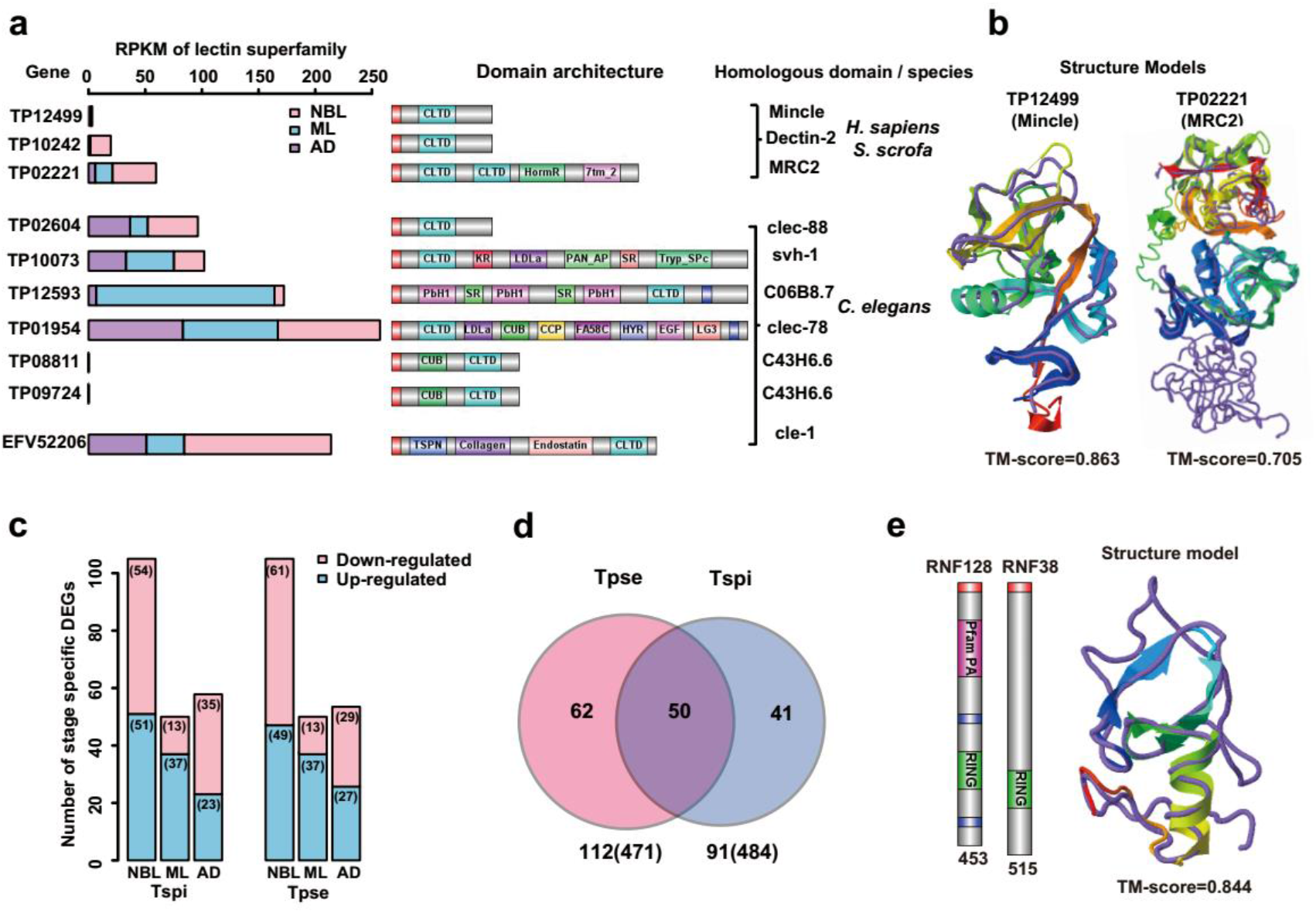
Stage-specific expressed E/S genes form the basis of varied parasitism between *T. pseudospiralis* and *T. spiralis*. **(a)** RPKM (left), domain architecture (middle) and homologous domain sequences (right) in mammalian host of C-type lectin superfamily. **(b)** Structural model of the C-type lectin domains of TP12499 and TP02221 are shown in cartoon, while the structural analog in the PDB library is displayed using backbone trace and colored in purple (as identified by TM-align). The TM-score value scales the structural similarity of the two structures. **(c)** The histogram shows numbers of stage-specifically differential expressed genes (DEGs) between the two species across three life stages. **(d)** Venn diagram of homologous SCOs between total SCOs identified in the two species. **(e)** Domain architectures and structural model of RNF128 and RNF38. The subgraph numbers indicate the protein length. Query structure is shown in cartoon, while the structural analog in the PDB library is displayed using backbone trace and colored in purple (as identified by TM-align). The TM-score value scales the structural similarity of the two structures.

In addition, we addressed the stage-specifically expressed genes between the two species and identified, 112 and 91 stage-specific single-copy orthologous (SCO) E/S genes in *T. pseudospiralis* and *T. spiralis*, respectively, among which 50 were homologous between the two genomes (Figs 5c and 5d); 34 out of the 50 genes were differentially expressed between the two species (Fisher test: *P* < 0.01) (Supplementary Fig. 19). Of note, 6 genes showed species-specific divergence in at least two stages of *T. pseudospiralis* (Supplementary Table 12), which were mostly reported as being involved in parasitism, as evident by enhanced parasite invasion and motility (Kostianovsky et al. 2007; Hashimoto et al. 2010; Helwig et al. 2013; Rausch and Hastings 2015; Zhao et al. 2017). The sequences of these genes were then further curated by Sanger sequencing, considering their potential significance in *Trichinella* parasitism (Supplementary Fig. 20). Among these genes, RNF128 expression was significantly up-regulated across three life stages in *T. pseudospiralis*. Domain analyses revealed that RNF128 had a zinc-finger (RING finger) motif at its C-terminal region and shared significant protein sequence and structure similarity with the human E3 ubiquitin-protein ligase, RNF38 (homology modeling TM-score=0.844) (Fig. 5e), which may specially bind host E2 enzyme and function in parasite-infected cells, to affect the stability or function of a number of host proteins, as supported by a previous study in *Trypanosoma cruzi* (Hashimoto et al. 2010). However, the feasibility of these genes in developing immunological assays needs further experimental evaluations.

## Discussion

Extensive efforts have been made to reveal the biological functions hidden in the repetitive portion of the genome (S and G 2006; Chalopin et al. 2015; EB et al. 2017; M et al. 2017). Despite of that, the *Trichinella spp*. represent an ideal model of studying the role of repetitive sequences in association with host adaptation. However, a large proportion of the published *T. pseudospiralis* genome was missing (Korhonen et al. 2016), as only 45 Mb of the genome was assembled, which is much less than 70 Mb of the genome size of *Trichinella spp.* (Zarlenga et al. 2009; Mitreva et al. 2011). Therefore, to compare genomes between encapsulated species (*T. spiralis*) and non-encapsulated species (*T. pseudospiralis*), which induce distinct differences in parasitism, pathology and immune response, high-quality *T. pseudospiralis* reference genome is needed.

In the present study, we have assembled a new *T. pseudospiralis* reference genome with large amounts of repeat sequences assembled, and annotated the genome and epigenome accordingly. Comparative analyses revealed that the most significant differences between the two genomes lay within the repeat content, as 27.7 Mb and 13.1 Mb repeat sequences were identified in *T. pseudospiralis* and *T. spiralis* genomes, respectively. Stepwise buildup of repetitive sequences not only has implications on genome stability, but also is considered as the driving force of epigenetic regulation through affecting DNA methylation patterns (Jurka et al. 2007). Thereby, the two *Trichinella* parasites differ significantly from each other both at genomic and epigenomic level, forming the foundations for differential parasitism between the two *Trichinella* species.

We further demonstrated that transposon elements could potentially facilitate different types of segmental duplications based on their density and distribution, suggesting for its pivotal role in gene duplication in the two *Trichinella* species. A variety of mechanisms could be involved in the generation of gene duplication, and the widely recognized one was likely mediated by repetitive sequences (Mendivil Ramos and Ferrier 2012). In clonal raider ant, approx. 1.5× higher transposon density than both the entire genome and other gene-dense regions of the genome were observed in odorant receptor (OR) loci, potentially facilitating tandem duplication by unequal crossing over (McKenzie and Kronauer 2018). However, within the *Drosophila melanogaster* subgroup, excess of repetitive sequences around flanking regions might mediate non-homologous recombination for generation of dispersed duplicate genes (Yang et al. 2008). Similarly, we observed higher TE density in *DNaseII* loci in *T. spiralis*, which may account for vast amounts of tandem duplication events by non-homologous end-joining. Such an extensive repertoire of *DNaseII* in *T. spiralis* is of particular importance for the intracellular parasitic mechanisms, especially for the nurse cell formation that is different from *T. pseudospiralis* in activation, proliferation and fusion of satellite cell, re-differentiation and transformation of infected muscle cells (Wu et al. 2001a). In contrast to the *DNaseII* family, we observed higher TE density around the expanded GSTs family in *T. pseudospiralis*, which may act as a route to interspersed duplication via nonallelic homologous recombination mediated by repetitive sequences, similar with previous studies in *D. melanogaster* (Yang et al. 2008; Mendivil Ramos and Ferrier 2012). The expanded GSTs endowed *T. pseudospiralis* enhanced invasion and migration in order to adapt to its non-encapsulated phenotype. However, such duplication also have immediate adverse effect on species fitness as it could lead to gene dosage imbalance. Our another important finding was that reduction in post-duplication gene expression is mediated through DNA methylation by inhibiting transcription initiation and elongation, further demonstrating close interplay between TEs, gene duplication and DNA methylation, as previously proved (Chang and Liao 2012; McKenzie and Kronauer 2018).

ML of Trichinella can survive for long term in muscle cells, and ML can induce or reform the infected muscle cells into a niche to nurse them for survival, which is related with function of E/S products (Liu et al. 2012). We present the landscape of transcriptomic changes between the two species and suggest aspects of differential immune system that might be associated with differential pathological characteristics. Several sets of E/S proteins that function at the host-parasite interface showed differential expression between the two species (for example, C-type lectins, RNF128, and other parasitic genes like PPT), which may probably be associated with the establishment of enhanced parasite invasion and motility in *T. pseudospiralis* (Hashimoto et al. 2010; Child et al. 2013; Guo et al. 2018). Our analysis demonstrated that *T. pseudospiralis* showed enhanced ability of immune evasion through higher expression of secreted C-type lectins and PPT. Moreover, we also observed the secreted proteins of RNF128 possessed E3 activity, which may function as an E3 ligase to a number of host proteins, as supported by a previous study in *Trypanosoma cruzi* (Hashimoto et al. 2010). These genes may be of particular importance in *T. pseudospiralis* survival and/or adaption to the non-encapsulated phenotype, which represents a novel finding with respect to new parasite immune-evasion strategy in *Trichinella* spp..

In summary, our present study published a high-quality reference genome of non-encapsulated *T. pseudospiralis*. We describes the first comprehensive comparative genomic and epigenomic study for understanding transposon element and DNA methylation variation in *Trichinella* and their roles on parasitism-related gene family expansion, potentially resulting in differential parasitology and immunology between the two *Trichinella* species. In addition, we also found a great number of novel genes that are related to the unelucidated puzzle of differences in parasitism, pathology and immune response. Through these characteristic comparisons, we provide new insight into the complex, distinct and differential parasitism in the intracellular nematodes, which may provided a foundation for further elucidation of mechanism of nurse cell formation and immunoevasion as well as identification of phamarcological and diagnostic targets of trichinellosis. Our approaches used here will be applicable to other parasitic nematodes with public health and food safety importance.

## Methods

### Generation and maintenance of *T. pseudospiralis* inbred strains

*Trichinella pseudospiralis* (ISS13), genotyped and proved by OIE Collaborating Center on Foodborne Parasites in Asian-Pacific Region, was preserved by serial passage in BALB/c mice. Recombinant inbred *T. pseudospiralis* strains were used in this study. To generate genetically homogeneous strains, sister-brother matings were performed for many consecutive generations. Typically, the mice were fed with a single pair of male and female ML selected from offspring obtained from the single pair of parents. A novel inbred *T. pseudospiralis* strain was established over >20 continuous generations after repeating the above operation for at least 20 times. ML were recovered from infected mice 35 dpi by artificial digestion (magnetic stirrer method) according to the OIE standard protocol. The Ad and NBL were isolated from the infected mice intestines, as described previously (Appleton and Mcgregor 1985; Liu et al. 2011). The animals were treated in strict according to the guidelines of the National Institutes of Health (publication no. 85-23, revised 1996). All experimental protocol involving animals have been reviewed and approved by the Ethical Committee of the Jilin University affiliated with the Provincial Animal Health Committee, Jilin Province, China (Ethical Clearance number IZ-2009-08).

### DNA preparation and high-throughput sequencing

High molecular weight genomic DNA were isolated from freshly collected muscle larvae. Total DNA amounts were determined using a Qubit Fluorometer dsDNA HS Kit (Invitrogen), according to the manufacturer’s instructions. Genomic DNA integrity was verified by agarose gel electrophoresis and using a BioAnalyzer (2100, Agilent). A paired-end library was constructed with an average insert size of 200 bp following the instructions provided by Illumina. Sequencing was performed on the Illumina Hiseq 2500 platform. A 20-kb library was constructed for PacBio sequencing on a PacBio RS II platform (Pacific Biosciences). Both the Illumina sequencing reads and Pacbio SMRT sequencing data were used for the genome assembly.

### Genome assembly

FastQC (v0.11.7) (https://github.com/s-andrews/FastQC) was used to assess the quality of raw sequencing reads. For Illumina reads, adapters and low-quality reads were filtered by Trimmomatic (v0.36) (Bolger et al. 2014) (LEADING:3, TRAILING:3, SLIDINGWINDOW:4:15, MINLEN:75), approx. 6.90 billion high-quality data were generated. For PacBio reads, low-quality subreads (error rate > 0.2), short subreads (< 5 kb), and duplicated reads were filtered to yield 9.93 billion of clean data. Illumina clean reads were assembled using Soapdenovo (v2.04) (Luo et al. 2012) with *k*-mer = 31. PacBio reads were assembled using Canu (v1.0) (Koren et al. 2017) algorithm, with the following parameters: error Rate = 0.01, genome Size = 70 Mb in addition to default parameters. Five mate-paired libraries were downloaded from NCBI (PRJNA257433) based on PE relationships. Pilon (v1.22) (Walker et al. 2014) algorithm was then used to correct the Canu assembly using recommended settings. Then, the two initially assembled genomes were subjected to a process of merging and scaffolding with long-insert mate-paired reads by SSPACE (v3.0) (Boetzer M 2014). Gap filling was performed by GapCloser (v1.12) (Luo et al. 2012) with the following parameters: GapCloser -a scafSeq -b gap_all.lib -o scafSeq.FG1 -t 10.

### Repetitive sequences annotation

Repeats were identified using a combination of *de-novo* and homology-based approaches. Two *de novo* software packages, LTR_FINDER (v1.07) (Xu and Wang 2007) and RepeatModeler (v1.0.11) (http://www.repeatmasker.org/RepeatModeler/), were used. LTR_FINDER searches the entire genome for finding full-length long terminal repeat retrotransposons (LTRs), while RepeatModeler is comprised of two programs (RECON and RepeatScout), which employ complementary computational methods for identifying repeat element boundaries and family relationships from sequence data. All repeat sequences with lengths > 100 bp and gap ‘N’ < 5% constituted the repeat element libraries. To further filter repeat sequences that are false positives, which are often genes or gene families containing transposable element domains, blast searches were conducted with the Nucleotide database in NCBI with an at least 50% identity over 50% length. Repeat sequences that meeting the threshold were removed from the libraries. We therefore constructed the repeat libraries that contained 766 sequences, of which 491 and 275 sequences derived from RepeatModeler and LTR_FINDER, respectively (Supplementary Table 8). Two software packages, including Tandem Repeats Finder (v4.09) and RepeatMasker (v4.0.7) (http://repeatmasker.org), were used in the homology-based prediction. Tandem repeats, which are two or more adjacent in DNA, were searched across the whole genome using the software package of Tandem Repeats Finder. RepeatMasker, which is comprised of two programs (RepeatMasker and RepeatProteinMask), was employed for DNA and protein level identification. TEs with 80% identity over 80% overlapping region were integrated together to construct a non-redundant repeat libraries. Transposon density of specific regions was calculated based on repeats belonging to known transposon classes. A detailed description of genome annotation is available at Supplementary Information.

### WGBS library construction and data processing

Two to five micrograms gDNA were sonicated to an approx. mean size range of 100-500 bp. After fragmentation, end-repair, addition of 3’ A bases and ligation of methylated cytosine PE adapters were performed, followed by bisulfite conversion of purified adapter-ligated DNA using an EZ DNA Methylation-Gold Kit™ (ZYMO Research, Irvine, CA, USA), according to the manufacturer’s instructions. Size selection was achieved by PAGE gel and yielded DNA fragments of 250-450 bp from bisulfite conversion of purified adapter-ligated DNA. The converted DNA was then purified with the QIAquick Gel Extraction Kit, followed by PCR enrichment using JumpStart™ Taq DNA Polymerase for eleven cycles with Illumina PE PCR primers. Methylated-adapter ligated to unmethylated lambda-phage DNA (Promega, Madison, WI, USA) was used as an internal control for assessing the bisulfite conversion ratio. Libraries were sequenced on Illumina HiSeq 2500 platform. Raw sequencing data were filtered for adapter contamination by cutadapt (Martin 2011), parsed through quality filtration (quality cut off value = 5, low-quality rate < 0.5, ‘N’ rate < 0.1) and the trimmed reads shorter than 50 bp were discarded. Clean reads from each library were mapped to the updated genome, using BSMAP (v2.73) (Xi and Li 2009). The methylation level of specific cytosine residues was estimated from the fraction of methylated sequence reads at that site (≥ 5x read depth).

### Transcriptome sequencing and differential expression

Total RNA from *T. pseudospiralis* (Ad, NBL, and ML) was purified using Trizol reagent (Invitrogen, CA, USA), according to the manufacturer’s instructions. RNA was dissolved in diethylpyrocarbonate (DEPC)-treated water and treated with DNase I (Invitrogen, CA, USA). The quantity and quality of the RNA were tested by ultraviolet-Vis spectrophotometry using a NanoDrop 2000 (Thermo Scientific CA, USA). Pair-end RNA-seq libraries were constructed, following Illumina’s protocols, for the three life stages of *T. pseudospiralis* and sequenced on an Illumina HiSeq 2500 platform. Raw sequencing data were filtered for adapter contamination by cutadapt (http://code.google.com/p/cutadapt/), parsed through quality filtration using the program Trimmomatic (v0.36) (Bolger et al. 2014) (Phred ≥ 20). TopHat (v2.0.12) (Kim et al. 2013) and followed by Cufflinks (v2.2.1) (Trapnell et al. 2012) to assemble the genome-referenced transcripts with default settings. Trinity (r20140717) (Haas et al. 2013) was used to conduct the de novo assembly. Gene expression analysis was measured using the reads per kb per million mapped reads and was calculated using Cufflinks. Differentially expressed genes were assessed with DESeq2 (http://www.bioconductor.org/packages/release/bioc/html/DESeq2.html).

### Gene family expansion and contraction analyses

OrthoMCL (Fischer et al. 2011) was used to confirm the genes that were orthologous among the following species: *T. pseudospiralis*, *T. spiralis*, *B. malayi*, *C. elegans*, *M. incognita*, *D. melanogaster* (outgroup). Single copy orthologous groups were extracted for phylogenetic tree construction using MrBayes (v3.2.2) (Ronquist et al. 2012) after aligning the family members with MUSCLE(Chojnacki et al. 2017). The optimal substitution models for amino acid and CDS sequences were estimated by ProtTest (v3.4.2) (Abascal et al. 2005) and ModelTest (v0.1.0) (Posada 2006), respectively. The divergence time for the six species using SCO gene families was estimated using the program MCMCTREE implemented in the PAML package(Yang 1997). The expansion or contraction events were determined using CAFÉ based on the comparison of orthologous gene family size differences under probabilistic graphical models (De Bie et al. 2006).

### Domain feature prediction and protein structural modeling

Conserved domains and transmembrane regions were identified using InterProScan and SMART (http://smart.embl-heidelberg.de/smart/set_mode.cgi). DOG (Domain Graph, version 2.0) (Emig et al. 2010) was used to plot the domain architecture. I-TASSER (https://zhanglab.ccmb.med.umich.edu/I-TASSER/), which is a hierarchical approach to protein structure and function prediction, was used to predict protein 3D-structure from the PDB by multiple threading approach LOMETS (Local Meta-Threading Server). Structural alignment was performed by TM-align (https://zhanglab.ccmb.med.umich.edu/TM-align/), which is an algorithm for sequence independent protein structure comparisons. The representative models were visualized by PyMOL (https://pymol.org/2/).

## Data availability

Raw genome sequencing data were deposited in the National Center for Biotechnology Information with the following accession number: SAMN08905168 under project PRJNA451013; SRP140458 for transcriptome and WGBS data; The genome sequence has also been deposited at DDBJ/ENA/GenBank (accession number QAWF00000000).

## Acknowledgments

We thank William J Lucas for help in editing the manuscript for grammar and writing style. We thank J. Ruan, P Cui and Q. Lin for helpful guidance on bioinformatic analysis. This study was supported by the National Natural Science Foundation of China (31520103916, 31872467), the National Key Research and Development Program of China (2017YFD0501302, 2017YFC1601206, 2016YFD0500707, 2018YFC1602504), Guangdong Innovative and Entrepreneurial Research Team Program (No.2014ZT05S123), Program for JLU Science and Technology Innovative Research Team, The Agricultural Science and Technology Innovation Program and The Elite Young Scientists Program of CAAS.

## Author contributions

M. Liu and F. Gao conceived the project. X. Liu and B. Tang undertook sample collection. X. Bai and X. Wang performed DNA extraction, and X. Liu and Y. Yang performed RNA extraction experiments. R. Qin and X. Yu undertook WGBS and RNA-seq library construction. Y. Feng carried out bioinformatic analysis. Y. Feng, X. Liu and F. Gao wrote the manuscript. All authors have read and approved the manuscript for publication.

## References

Abascal F, Zardoya R, Posada D. 2005. ProtTest: selection of best-fit models of protein evolution. Bioinformatics 21(9): 2104–2105.

Ae C, Nm A, M K, Aj S, Rm P. 1986. Repetitive DNA as a tool for the identification and comparison of nematode variants: application to Trichinella isolates. Molecular and biochemical parasitology 21(2): 113–120.

Al-Mulla Hummadi YM, Al-Bashir NM, Najim RA. 2006. Leishmania major and Leishmania tropica: II. Effect of an immunomodulator, S(2) complex on the enzymes of the parasites. Experimental parasitology 112(2): 85–91.

Appleton JA, Mcgregor DD. 1985. Life-phase specific induction and expression of rapid expulsion in rats suckling Trichinella spiralis-infected dams. Immunology 55(2): 225–232.

Boetzer M PW. 2014. SSPACE-LongRead: scaffolding bacterial draft genomes using long read sequence information. BMC Bioinformatics 15(1).

Bolger AM, Lohse M, Usadel B. 2014. Trimmomatic: a flexible trimmer for Illumina sequence data. Bioinformatics 30(15): 2114–2120.

Boonmars T, Wu Z, Nagano I, Takahashi Y. 2005. Trichinella pseudospiralis infection is characterized by more continuous and diffuse myopathy than T. spiralis infection. Parasitology research 97(1): 13–20.

Borgognone A, Castanera R, Morselli M, Lopez-Varas L, Rubbi L, Pisabarro AG, Pellegrini M, Ramirez L. 2018. Transposon-associated epigenetic silencing during Pleurotus ostreatus life cycle. DNA research : an international journal for rapid publication of reports on genes and genomes 25(5): 451–464.

Chalopin D, Naville M, Plard F, Galiana D, Volff JN. 2015. Comparative analysis of transposable elements highlights mobilome diversity and evolution in vertebrates. Genome biology and evolution 7(2): 567–580.

Chang AY, Liao BY. 2012. DNA methylation rebalances gene dosage after mammalian gene duplications. Molecular biology and evolution 29(1): 133–144.

Child MA, Hall CI, Beck JR, Ofori LO, Albrow VE, Garland M, Bowyer PW, Bradley PJ, Powers JC, Boothroyd JC et al. 2013. Small-molecule inhibition of a depalmitoylase enhances Toxoplasma host-cell invasion. Nature chemical biology 9(10): 651–656.

Chojnacki S, Cowley A, Lee J, Foix A, Lopez R. 2017. Programmatic access to bioinformatics tools from EMBL-EBI update: 2017. Nucleic acids research 45(W1): W550–W553.

De Bie T, Cristianini N, Demuth JP, Hahn MW. 2006. CAFE: a computational tool for the study of gene family evolution. Bioinformatics 22(10): 1269–1271.

Dermitzakis ET, Clark AG. 2001. Differential Selection After Duplication in Mammalian Developmental Genes. Molecular biology and evolution 18(4): 557–562.

Ds Z, G LR. 2000. Molecular and biochemical methods for parasite differentiation within the genus Trichinella. Veterinary parasitology 93(null): 279–292.

Eb C, Nc E, C F. 2017. Regulatory activities of transposable elements: from conflicts to benefits. Nature reviews Genetics 18(2): 71–86.

Emig D, Salomonis N, Baumbach J, Lengauer T, Conklin BR, Albrecht M. 2010. AltAnalyze and DomainGraph: analyzing and visualizing exon expression data. Nucleic acids research 38(Web Server issue): W755–762.

Fischer S, Brunk BP, Chen F, Gao X, Harb OS, Iodice JB, Shanmugam D, Roos DS, Stoeckert CJ, Jr. 2011. Using OrthoMCL to assign proteins to OrthoMCL-DB groups or to cluster proteomes into new ortholog groups. Current protocols in bioinformatics Chapter 6: Unit 6 12 11–19.

Gao F, Liu X, Wu XP, Wang XL, Gong D, Lu H, Xia Y, Song Y, Wang J, Du J et al. 2012. Differential DNA methylation in discrete developmental stages of the parasitic nematode Trichinella spiralis. Genome biology 13(10): R100.

Gottstein B, Pozio E, Nockler K. 2009. Epidemiology, diagnosis, treatment, and control of trichinellosis. Clinical microbiology reviews 22(1): 127–145, Table of Contents.

Guo Y, Chang Q, Cheng L, Xiong S, Jia X, Lin X, Zhao X. 2018. C-Type Lectin Receptor CD23 Is Required for Host Defense against Candida albicans and Aspergillus fumigatus Infection. Journal of immunology 201(8): 2427–2440.

Haas BJ, Papanicolaou A, Yassour M, Grabherr M, Blood PD, Bowden J, Couger MB, Eccles D, Li B, Lieber M et al. 2013. De novo transcript sequence reconstruction from RNA-seq using the Trinity platform for reference generation and analysis. Nature protocols 8(8): 1494–1512.

Hashimoto M, Murata E, Aoki T. 2010. Secretory protein with RING finger domain (SPRING) specific to Trypanosoma cruzi is directed, as a ubiquitin ligase related protein, to the nucleus of host cells. Cellular microbiology 12(1): 19–30.

Helwig M, Hoshino A, Berridge C, Lee SN, Lorenzen N, Otzen DE, Eriksen JL, Lindberg I. 2013. The neuroendocrine protein 7B2 suppresses the aggregation of neurodegenerative disease-related proteins. The Journal of biological chemistry 288(2): 1114–1124.

International Helminth Genomes C. 2019. Comparative genomics of the major parasitic worms. Nature genetics 51(1): 163–174.

Jurka J, Kapitonov VV, Kohany O, Jurka MV. 2007. Repetitive sequences in complex genomes: structure and evolution. Annu Rev Genomics Hum Genet 8: 241–259.

Kaessmann H. 2010. Origins, evolution, and phenotypic impact of new genes. Genome research 20(10): 1313–1326.

Kim D, Pertea G, Trapnell C, Pimentel H, Kelley R, Salzberg SL. 2013. TopHat2: accurate alignment of transcriptomes in the presence of insertions, deletions and gene fusions. Genome biology 14: R36.

Koren S, Walenz BP, Berlin K, Miller JR, Bergman NH, Phillippy AM. 2017. Canu: scalable and accurate long-read assembly via adaptive k-mer weighting and repeat separation. Genome research 27(5): 722–736.

Korhonen PK, Pozio E, La Rosa G, Chang BC, Koehler AV, Hoberg EP, Boag PR, Tan P, Jex AR, Hofmann A et al. 2016. Phylogenomic and biogeographic reconstruction of the Trichinella complex. Nature communications 7: 10513.

Kostianovsky AM, Maier LM, Baecher-Allan C, Anderson AC, Anderson DE. 2007. Up-Regulation of Gene Related to Anergy in Lymphocytes Is Associated with Notch-Mediated Human T Cell Suppression. The Journal of Immunology 178(10): 6158–6163.

Li LG, Wang ZQ, Liu RD, Yang X, Liu LN, Sun GG, Jiang P, Zhang X, Zhang GY, Cui J. 2015. Trichinella spiralis: low vaccine potential of glutathione S-transferase against infections in mice. Acta tropica 146: 25–32.

Liu CY, Ren HN, Song YY, Sun GG, Liu RD, Jiang P, Long SR, Zhang X, Wang ZQ, Cui J. 2018. Characterization of a putative glutathione S-transferase of the parasitic nematode Trichinella spiralis. Experimental parasitology 187: 59–66.

Liu X, Song Y, Jiang N, Wang J, Tang B, Lu H, Peng S, Chang Z, Tang Y, Yin J et al. 2012. Global gene expression analysis of the zoonotic parasite Trichinella spiralis revealed novel genes in host parasite interaction. PLoS neglected tropical diseases 6(8): e1794.

Liu X, Song Y, Lu H, Tang B, Piao X, Hou N, Peng S, Jiang N, Yin J, Liu M et al. 2011. Transcriptome of small regulatory RNAs in the development of the zoonotic parasite Trichinella spiralis. PloS one 6(11): e26448.

Loukas A. 2000. Helminth C-type lectins and host-parasite interactions. Parasitol Today 16(8): 333–339.

Lu X, Nagata M, Yamasaki S. 2018. Mincle: 20 years of a versatile sensor of insults. Int Immunol 30(6): 233–239.

Luo R, Liu B, Xie Y, Li Z, Huang W, Yuan J, He G, Chen Y, Pan Q, Liu Y et al. 2012. SOAPdenovo2: an empirically improved memory-efficient short-read de novo assembler. Gigascience 1(1): 2047-2217X–2041-2018.

M T, Y P, M H-B, K A, K M, M C, Gh P, Vj L, Cd B. 2017. Transposable elements are the primary source of novelty in primate gene regulation. Genome research 27(10): 1623–1633.

Martin M. 2011. Cutadapt removes adapter sequences from high-throughput sequencing reads. Embnet Journal 17(1).

McKenzie SK, Kronauer DJC. 2018. The genomic architecture and molecular evolution of ant odorant receptors. Genome research 28(11): 1757–1765.

Mendivil Ramos O, Ferrier DE. 2012. Mechanisms of Gene Duplication and Translocation and Progress towards Understanding Their Relative Contributions to Animal Genome Evolution. International journal of evolutionary biology 2012: 846421.

Mitreva M, Jasmer DP, Zarlenga DS, Wang Z, Abubucker S, Martin J, Taylor CM, Yin Y, Fulton L, Minx P et al. 2011. The draft genome of the parasitic nematode Trichinella spiralis. Nature genetics 43(3): 228–235.

Mr C, E R, Rm P, Pr B, T G. 1994. The differentiation of parasitic nematodes using random amplified polymorphic DNA. Journal of helminthology 68(2): 109–113.

Nagano I, Wu Z, Takahashi Y. 2009. Functional genes and proteins of Trichinella spp. Parasitology research 104(2): 197–207.

Oliver KR, Greene WK. 2009. Transposable elements: powerful facilitators of evolution. BioEssays 31(7): 703–714.

Posada D. 2006. ModelTest Server: a web-based tool for the statistical selection of models of nucleotide substitution online. Nucleic acids research 34(Web Server issue): W700–703.

Rausch MP, Hastings KT. 2015. Diverse cellular and organismal functions of the lysosomal thiol reductase GILT. Molecular immunology 68(2 Pt A): 124–128.

Robinson MW, Greig R, Beattie KA, Lamont DJ, Connolly B. 2007. Comparative analysis of the excretory-secretory proteome of the muscle larva of Trichinella pseudospiralis and Trichinella spiralis. International journal for parasitology 37(2): 139–148.

Ronquist F, Teslenko M, van der Mark P, Ayres DL, Darling A, Hohna S, Larget B, Liu L, Suchard MA, Huelsenbeck JP. 2012. MrBayes 3.2: efficient Bayesian phylogenetic inference and model choice across a large model space. Systematic biology 61(3): 539–542.

S T, G W. 2006. Differential evolution of repetitive sequences in Cryptosporidium parvum and Cryptosporidium hominis. Infection, genetics and evolution : journal of molecular epidemiology and evolutionary genetics in infectious diseases 6(2): 113–122.

Sarda S, Zeng J, Hunt BG, Yi SV. 2012. The evolution of invertebrate gene body methylation. Molecular biology and evolution 29(8): 1907–1916.

Schrader L, Kim JW, Ence D, Zimin A, Klein A, Wyschetzki K, Weichselgartner T, Kemena C, Stokl J, Schultner E et al. 2014. Transposable element islands facilitate adaptation to novel environments in an invasive species. Nature communications 5: 5495.

Trapnell C, Roberts A, Goff L, Pertea G, Kim D, Kelley DR, Pimentel H, Salzberg SL, Rinn JL, Pachter L. 2012. Differential gene and transcript expression analysis of RNA-seq experiments with TopHat and Cufflinks. Nature protocols 7(3): 562–578.

Tsagkogeorga G, Muller S, Dessimoz C, Rossiter SJ. 2017. Comparative genomics reveals contraction in olfactory receptor genes in bats. Scientific reports 7(1): 259.

Tyson JR, O’Neil NJ, Jain M, Olsen HE, Hieter P, Snutch TP. 2018. MinION-based long-read sequencing and assembly extends the Caenorhabditis elegans reference genome. Genome research 28(2): 266–274.

Walker B, Abeel T, Shea T, Priest M, Abouelliel A, Sakthikumar S, Cuomo C, Zeng Q, Wortman J, Young S et al. 2014. Pilon: An Integrated Tool for Comprehensive Microbial Variant Detection and Genome Assembly Improvement. PloS one 9.

Wang S, Wang S, Luo Y, Xiao L, Luo X, Gao S, Dou Y, Zhang H, Guo A, Meng Q et al. 2016. Comparative genomics reveals adaptive evolution of Asian tapeworm in switching to a new intermediate host. Nature communications 7: 12845.

Wu Z, Matsuo A, Nakada T, Nagano I, Takahashi Y. 2001a. Different response of satellite cells in the kinetics of myogenic regulatory factors and ultrastructural pathology after Trichinella spiralis and T. pseudospiralis infection. Parasitology 123(01).

Wu Z, Matsuo A, Nakada T, Nagano I, Takahashi Y. 2001b. Different response of satellite cells in the kinetics of myogenic regulatory factors and ultrastructural pathology after Trichinella spiralis and T. pseudospiralis infection. Parasitology 123(Pt 1): 85–94.

Xi Y, Li W. 2009. BSMAP: whole genome bisulfite sequence MAPping program. BMC bioinformatics 10: 232.

Xiao S, Xie D, Cao X, Yu P, Xing X, Chen CC, Musselman M, Xie M, West FD, Lewin HA et al. 2012. Comparative epigenomic annotation of regulatory DNA. Cell 149(6): 1381–1392.

Xu Z, Wang H. 2007. LTR_FINDER: an efficient tool for the prediction of full-length LTR retrotransposons. Nucleic acids research 35(Web Server issue): W265–268.

Yang S, Arguello JR, Li X, Ding Y, Zhou Q, Chen Y, Zhang Y, Zhao R, Brunet F, Peng L et al. 2008. Repetitive element-mediated recombination as a mechanism for new gene origination in Drosophila. PLoS genetics 4(1): e3.

Yang Z. 1997. PAML: A program package for phylogenetic analysis by maximum likelihood. Computer applications in the biosciences: CABIOS 13(5): 555–556.

Z W, I N, T N, Y T. 2002. Expression of excretory and secretory protein genes of Trichinella at muscle stage differs before and after cyst formation. Parasitology international 51(2): 155–161.

Zarlenga DS, Rosenthal B, Hoberg E, Mitreva M. 2009. Integrating genomics and phylogenetics in understanding the history of Trichinella species. Veterinary parasitology 159(3-4): 210–213.

Zarlenga DS, Rosenthal BM, La Rosa G, Pozio E, Hoberg EP. 2006. Post-Miocene expansion, colonization, and host switching drove speciation among extant nematodes of the archaic genus Trichinella. Proceedings of the National Academy of Sciences of the United States of America 103(19): 7354–7359.

Zhao L, Shao S, Chen Y, Sun X, Sun R, Huang J, Zhan B, Zhu X. 2017. Trichinella spiralis Calreticulin Binds Human Complement C1q As an Immune Evasion Strategy. Frontiers in immunology 8.

